# Mesoscale computational protocols for the design of highly cooperative bivalent macromolecules

**DOI:** 10.1101/743088

**Authors:** S. Saurabh, F. Piazza

## Abstract

The last decade has witnessed a swiftly increasing interest in the design and production of novel multivalent molecules as powerful alternatives for conventional antibodies in the fight against cancer and infectious diseases. However, while it is widely accepted that large-scale flexibility (10 − 100 nm) and free/constrained dynamics (100 ns − *µ*s) control the activity of such novel molecules, computational strategies at the mesoscale still lag behind experiments in optimizing the design of crucial features, such as the binding cooperativity (a.k.a. avidity).

In this study, we introduced different coarse-grained models of a polymer-linked, two-nanobody composite molecule, with the aim of laying down the physical bases of a thorough computational drug design protocol at the mesoscale. We show that the calculation of suitable potentials of mean force allows one to apprehend the nature, range and strength of the thermodynamic forces that govern the motion of free and wall-tethered molecules. Furthermore, we develop a simple computational strategy to quantify the encounter/dissociation dynamics between the free end of a wall-tethered molecule and the surface, at the roots of binding cooperativity. This procedure allows one to pinpoint the role of internal flexibility and weak non-specific interactions on the kinetic constants of the NB-wall encounter and dissociation. Finally, we quantify the role and weight of rare events, which are expected to play a major role in real-life situations, such as in the immune synapse, where the binding kinetics is likely dominated by fluctuations.

**SIGNIFICANCE:** Multivalent and multispecific molecules composed of polymer-linked nanobodies have gained interest as engineered alternatives to conventional antibodies. These therapeutic molecules have a larger reach due to their smaller size and promise substantial and tunable gains in avidity. This paper studies a model diabody to lay the bases of a multi-scale computational design of the structural and dynamical determinants of binding cooperativity, rooted in a blend of atomistic and coarse-grained MD simulations and concepts from statistical mechanics.

## INTRODUCTION

Single-domain antibodies, also known as *nanobodies* (NB) (1), are found naturally in camelids and represent an intriguing alternative to design and build novel multivalent and multi-specific immunotherapy agents for targeting tumors and viral infections (2). In addition to leading to an increase in affinity, coupling two (or more) different *binders* helps in marginalising the effect of mutation and polymorphism of the target. Linked anti-CD16 nanobodies, C21 and C28 (3), for example, have been shown to be effective against treating breast cancer involving a low HER2 expression, which is resistant to the therapeutic antibody trastuzumab (4).

As the conventional antibodies are large structures, they may not be effective when it comes to situations like a partially exposed tumor (5, 6). In addition, an uneven distribution of the anti-tumor antibody in the entire tumor region may lead to tumor regrowth (7). In such situations, therapeutic agents with smaller structures are desirable (8). A good solution is to use engineered structures composed of antigen-specific nanobodies linked with flexible linkers (9), e.g. realized via a polymer such as [Gly_4_Ser]_*n*_. Such structures will be small and will thus have better ability to penetrate into the tumor microenvironment and reinforce the formation of the immune synapse. Such structures are easy to produce and are more efficient when compared to the bulky conventional antibodies, and can be designed to be multivalent and multispecific (11, 12).

An important property of an antibody is the strength of bivalent binding that it demonstrates, a property known in immunology as *avidity*. Although it has no unique quantitative definition (let alone whether it is more appropriate to regard it as an equilibrium or a kinetic parameter), avidity can be thought of as the cooperative gain in affinity afforded by double binding. Notably, such measure is intimately connected to the internal flexibility of the multi-domain molecules and to the geometric configuration of binding epitopes (10). For example, virions employ various strategies to evade the action of antibodies of the immune system. Some virions, like HIV-1, have a very rapid rate of mutation. Experiments reveal that the enhanced antibody evasion capability of HIV-1 is based not just in its capability to mutate but is a combined effect of mutation along with the spike structure and low spike density (13–15). While, mutations reduce the affinity of the natural antibodies towards the target spikes, the spike structure and low density preclude intra- (i.e. multiple epitopes on the same target) and inter-spike cross-linking thus preventing bivalent binding and avidity (16, 17).

Experiments have demonstrated that linking two Fab domains of an antibody through an extended polymer like a DNA oligomer, leads to an increase in divalent binding, for an optimal linker length, and a resulting increase in the efficacy of the antibodies. Wu *et al.* (18) performed experiments on respiratory syncytial virus, which has a very high Env spike density, and showed that the affinity of low-affinity bivalent Fabs was 2−3 orders of magnitude higher as compared to their monovalent counterparts and the efficiency was not affected by mutations that increased the off-rates nearly 100-fold. The results show that multivalent structures made of polymer linked nanobodies would lead to higher degree of avidity and thus higher efficiency. Galimidi *et al.* (19) performed experiments on linked Env (HIV-1 envelope glycoproteins) binders and showed that linking can lead to an increase in potency by 2−3 orders of magnitude. Jähnichen et al. (20) developed two different single domain nanobodies that could bind to different sites on the extracellular domain of the CXCR4 coreceptor. They found that joining the two nanobodies with protein linkers resulted in a 27-fold increase in CXCR4 affinity. With further analysis they concluded that the effect is pure avidity resulting from the heterobivalent linking of the two nanobodies. Zhang *et al.* (21) developed multimers of nanobodies leading to an increase in affinity and several orders of magnitude decrease in the rates of dissociation. Yang *et al.* (22) performed dissociation rate calculations for the binding between a bivalent antibody and hapten ligands as a function of the ligand density. They found cooperative binding as the hapten density increased and bivalent binding set in. They could determine two different dissociation constants for the double-step antibody-hapten bnding process with one dissociation constant being 3-orders of magnitude larger than the other.

When one of the nanobodies in a two-NB construct binds to the receptor at its binding site (epitope), the other linked ligand spends more time in the vicinity, leading to a larger probability of the latter unit to bind to another similar or different epitope, on the same or on a facing surface, depending on whether the system is mono-specific or multi-specific. By hindering free diffusion of the ligands, linking can lead to an increase in rebinding events and strengthen the interaction between interfaces (23). Further, Bongrand *et al.* (24) showed that the fraction of divalent attachments between an antibody-coated microsphere and a mono- or divalent ligand-coated surface, that resisted a force of 30 pN for a minimum of 5 seconds, was ∼ 4 times higher than the number of monovalent attachments.

While the choice of the nanobody depends not only on its affinity towards the target epitope, but also on the nature of bond it forms with the epitope (25), the choice of linker would depend on its flexibility and the geometry of the epitope distribution/configurations. Experimental methods have been developed to create linkers of given stiffness and extension, out of a combination of proteins and peptides (26–29). Among the most common are the (Gly_4_Ser)_*n*_ linkers. Protein linkers have been accommodated into the hinge regions of natural antibodies, thus enabling intra-spike linking to viral receptors (28). Other biocompatible polymers like PEG are also good candidates to be used to link the nanobodies. The properties of the linker are very important in determining the degree of avidity. Depending on the epitope density on a tumor cell or the distribution of the antibody binding sites on the viral envelope, a linker that is either too flexible or too stiff can lead to under-performance.

With the improvements in computational resources and speed, molecular dynamics (MD) simulations in recent days have been playing an important role in fields like drug discovery (30) and also in unraveling fundamental mechanisms involving very large biological complexes, such as chromatin (31). MD simulations can be important in determining the optimal properties of the linkers that would lead to an efficient multivalent binding for a particular target. In addition, simulations of a group of linked nanobodies can give important insights into their epitope-binding kinetics as a function of the linker structural properties and other important parameters, like the paratope-epitope binding energy. While the engineered nanobody-linker-nanobody systems are much smaller as compared to the conventional antibodies, simulating a significantly large group of them in atomistic detail would be computationally expensive and cannot be done routinely. Thus, some degree of coarse-graining is important to perform kinetics analysis using MD simulation as a tool.

Keeping the above discussion in mind, here we perform MD simulation of a coarse-grained system consisting of two nanobodies connected by a linker (referred to as a diabody) and study its structural and dynamical properties. We use different levels of coarse graining schemes to represent the diabody. In the simplest representation, we perform simulation of two extended (rigid) spheres connected by a flexible bead-string linker (see Fig. 1). Similar model has been used previously to study the dynamics of a polymer tethered to a surface with a big bead at the free end (32). In addition to this, with a future aim to study the dynamics of diabodies in the presence of their target receptors (such as HER2), where a nanobody represented by a hard sphere will be incapable of representing the important features of the diabody-target interaction properly, we perform a finer coarse-graining of the nanobody (see Fig. 1). In this scheme, we represent the nanobody using the shape-based coarse-graining (SBCG) scheme developed by Schulten *et al.* (33). Using Umbrella Sampling (US) simulations we calculate the free energy profiles on which various conformations of diabodies tethered to a surface lie. From the MD trajectories we calculate the flight and residence times of the free end of the tethered diabody in a region close to the tethering wall. Comparing the results from the two different models we demonstrate how the coarse-graining scheme could affect the results and, notably, we highlight the role of the “shape” in the wall-domain dynamics.

**Figure 1:**
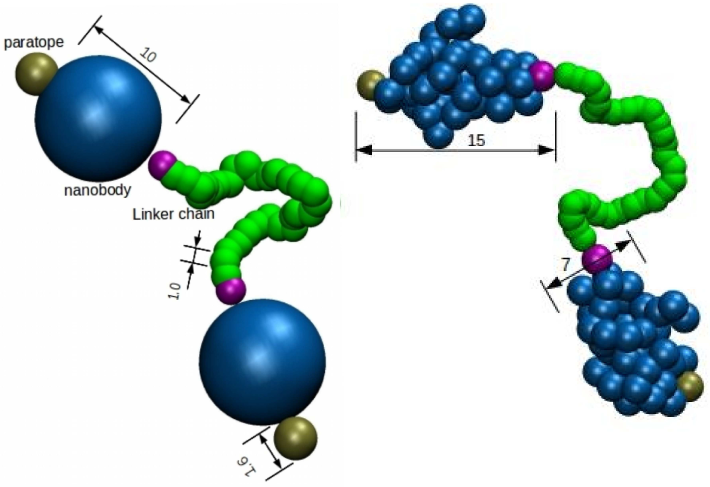
The two coarse-grained models of diabody studied in this work. Two nanobodies are linked through a polymer. The nanobody beads are represented in blue an the polymer in green. The maroon beads are the connector beads, which connect the nanobody to the polymer, and paratopes, respectively. The model on the left (named SPH) uses single spherical beads to represent the nanobodies. The model on the right (named SBCG) uses a shape-based coarse-graining algorithm for the nanobodies.

The paper is organize as follows. In section 1 we provide the details of our computational methods, models and simulations. In section 2, we describe and comment our results, mainly concerning the different potentials of mean force and the detailed analyses of the encounter and dissociation kinetics of the free NB of a wall-tethered molecule. In section 3 we wrap up our discussion and provide some final comments on this work and its perspectives.

## METHODS

### Coarse-graining schemes and simulations

All the data reported in this work were generated from Langevin dynamics simulations performed with LAMMPS (42) in the Lennard-Jones (LJ) unit system, with the unit of length being *σ* = 3.5 Å (the size of one monomer of the linker) and the unit of energy being *ϵ* = 100 K. All the simulations were conducted via Langevin dynamics in the overdamped regime.

In this work, we considered two different coarse-grained representations of a two-NB molecule joined by a flexible linker. In the first approach, the NBs were modeled as rigid spheres of radius 10 (in units of linker monomers), decorated with two smaller fixed spheres at two diametrically opposite ends, representing the NB-linker connecting unit and the paratope, respectively (see Fig.1). The paratope bead has diameter 1.6 and the connector bead diameter 1. A bead-spring polymer bridges the two connector beads. The beads constituting the polymer linker have unit diameter. This model will be referred to in the following as the SPH model.

The second model was meant to reproduce the *shape* and large-scale flexibility of the NBs. For that we used the shape-based coarse-graining scheme developed in Schulten’s group (33). This procedure requires a trajectory from an atomistic equilibrium MD simulation to be sampled and fed as an input. The crystal structure with pdb id: 1qd0 (34) was used as the starting structure for the MD simulation to generate the input structure. This is a camelid heavy chain variable (VHH) domain, in complex with a RR6 dye dimer.

The dye was removed from the complex and the remaining protein was solvated in TIP3P water with a 20 Å buffer, leading to a system size of 40298 atoms, with 13284 water molecules. The solvated system was then neutralized by adding 5 Cl^−^ ions to generate the starting configuration for the MD simulation (see Fig.2). The system was minimized for 10000 steps using the conjugate gradient method. During minimization, all the atoms in the protein were constrained to their starting positions. This allowed water molecules to re-organize and eliminate unfavorable contacts with the protein. After minimization, the atoms were assigned velocities generated from a Maxwell distribution at 300 K. The particle mesh Ewald (PME) method with a real space cut-off of 12 Å was used to estimate the energy component from the long-range electrostatic interaction. The system was simulated for 100 ns in the NPT ensemble. A Langevin thermostat was used (35) with a temperature coupling constant of 5 ps^−1^, while the pressure was regulated using the Langevin barostat with a pressure coupling constant of 50 ps^−1^. The simulation was performed using NAMD (36) and the CHARMM 27 (37) force field was used to describe the protein. The MD trajectories were visualized using VMD (38).

**Figure 2:**
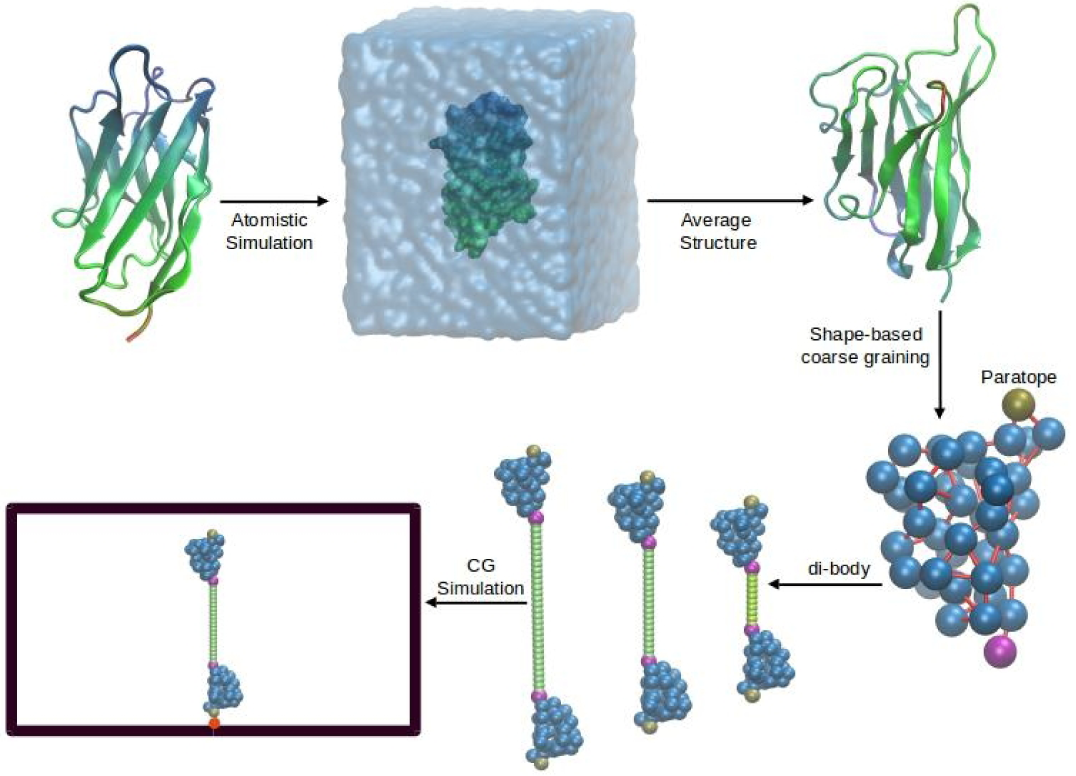
The coarse-graining procedure. The crystal structure (PDB id. 1qd0) is used as the starting configuration for the explicit solvent atomistic simulation. Snapshots from the last 10 ns of the trajectory are used to generate an average structure. The average structure is then coarse-grained using the shape-based coarse-graining method. A pair of the coarse-grained structures are connected by a linker (10-, 20- and 30-mer), enclosed in a box with *X* -*Y* periodicity, aligned along the *Z*-direction and tethered to the box base, thus generating the starting configuration for CG simulations.

An average structure was generated from the snapshots belonging to the last 10 ns of the MD trajectory (see Fig. 2). The average structure of the nanobody was used to generate a shape-based coarse grained (SBCG) model using the procedure formulated by Schulten *et al.* (33). This scheme uses topology-conserving algorithm developed for neural networks to generate a coarse-grained representation that reproduces the shape of the protein. In a trade-off between the system size and a good representation of the protein shape, we used 40 beads to represent the 126 residue protein. To generate the system to be simulated two of the CG proteins were connected by a polymer linker with monomer diameter 1, similar to the one used in the SPH model (see Fig.1). It is to be noted that the diameter of the spherical bead that represents the nanobody in the SPH model is nearly equal to the geometric average of the major and minor axes of the roughly spheroidal SBCG nanobody. In this sense the two models are equivalent and comparable.

### Interaction parameters

In general, the total interaction potential of our coarse-grained models had bond, angle and van der Waals (vdW) terms, given by

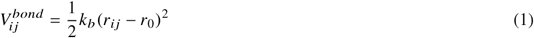

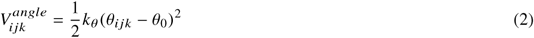

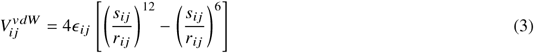

where *k*_*b*_ and *k*_*θ*_ are the bond force constant and the angle bending energy, respectively, *r*_0_ and *θ*_0_ are the equilibrium bond length and angle, respectively, *ϵ* _*ij*_ is the LJ interaction energy and *s* is the inter-bead distance at which the LJ potential becomes repulsive (referred to as *repulsive* length in the following), which depends on the combined radii of the interacting beads.

#### Interaction parameters for the SBCG nanobodies and the linker

All the CG beads were kept neutral. The connectivity and spring constants for the bonds between the beads were used as generated by the SBCG scheme. It is to be noted that the bonds are not set by a distance-based cut-off scheme, but are in accordance with the bonds present in the atomistic system. This helps in maintaining the flexibility of the nanobody and providing the required flexibility to the loop regions, which may play a defining role in the kinetics of the diabody. Repulsive vdW interactions among the beads were also introduced to ensure that the shape is maintained during the course of the simulation. The vdW radii of the protein beads were generated by comparing the masses of the protein beads to that of the PEG monomer. The repulsion between the constituent beads of the nanobody was represented by a Weeks-Chandler-Andersen potential (WCA) (39), i.e. a shifted LJ potential cut off at the minimum, i.e. r_*cut*_ = 2^1/6^ *σ*. The angle parameters for the nanobody beads were used as generated by the SBCG scheme.

The linker is represented as a freely-jointed chain with two-body harmonic bond-stretching and three-body harmonic bond-bending potentials. The masses and diameters of the polymer beads were set to 1. The vdW interaction between the bead pairs was set to purely repulsive as represented by a WCA potential. The force constant of the (stiff) bond-stretching potential was set to 54 N/m, which corresponds to an average fluctuation of the bonds at room temperature of about 2 % of the equilibrium length. In order to estimate the appropriate value of the bending rigidity *k*_*θ*_, it is expedient to refer to the calculation of the persistence length *t*_*p*_ for the freely jointed chain with angle-bending interactions in the hypothesis of zero correlation between bending and torsion degrees of freedom. Referring to published data for PEG (41), we fixed *k*_*θ*_ = 1.8 *k*_*B*_*T* as the bending coefficient for the linker (see (40) for a detailed discussion).

#### Interaction of the beads with the box walls

The walls of the simulation box in the *X* and *Y* directions had periodic boundary condition, while the *Z* walls were fixed. The Z walls interact with the beads via LJ (12-6) interaction given by

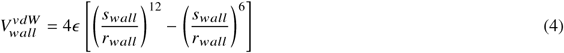

Here *r*_*wall*_ is the distance of the center of any bead from the wall. The wall interaction parameters, *s*_*wall*_, *r*_*cutwall*_ and *ϵ* define the nature of interaction between different beads and the wall. For the purely repulsive wall, *r*_*cutwall*_ = 2^1/6^ *s*_*wall*_, while for the attractive wall it equals 2.5 *s*_*wall*_ (see also Table 1). While the interaction of the nanobodies and paratopes with the *z* walls was set to be either repulsive or attractive in different simulations, the linker and connector beads always had repulsive interaction with the walls.

**Table 1:**
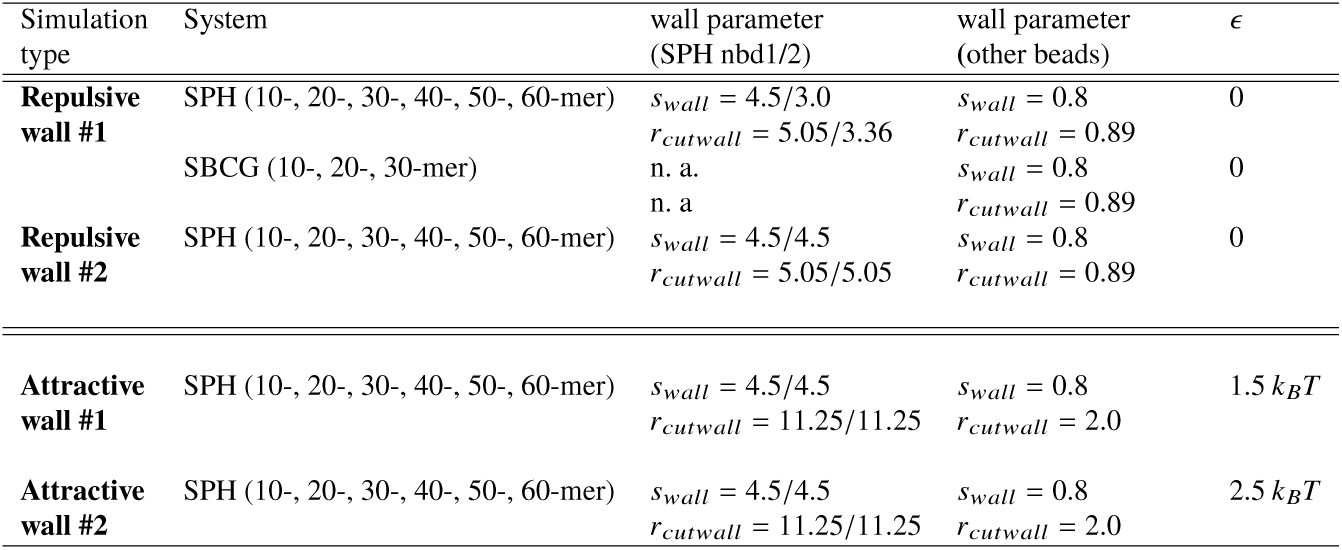
Simulation details. Four sets of simulations were performed for calculating the flight/residence times, corresponding to different choices of the parameters describing the interaction between CG beads forming the SBCG NBs and the wall.

### Preparing the systems for simulation

The first set of simulations reported involve the calculation of potentials of mean force (PMF) for the diabodies as a function of various reaction coordinates. To perform the PMF calculations, the diabodies were enclosed in cuboidal boxes which were periodic in the *X* and *Y* directions, while the *Z* walls were fixed and repulsive. The diabodies were tethered to the lower *Z* wall by imposing an attractive LJ interaction between the paratope of one of the nanobodies and a fixed epitope bead attached to the lower *z*-wall. The simulations were performed for linker lengths of 10, 20, 30, 40 and 50 monomers for the SPH system and 10, 20 and 30 monomers for the SBCG system.

The second kind of simulations were performed on a collection of diabodies to calculate dynamical parameters. *N* = 25 diabodies were tethered to the lower *Z*-wall of a cuboidal box. The tethering points were arranged in a 5 × 5 lattice (see Fig. 2 of supplementary information (SI) (40)). Again, the *x* and *y* directions had periodic boundary conditions, while the lower *z* wall was fixed and either perfectly reflecting or attractive. The linker lengths for different systems were similar to that considered for the PMF calculation, with an additional linker length of 60-mer for the SPH system.

In the rest of the article, the free nanobody is referred to as nbd-1 while the nanobody tethered to the wall is referred to as nbd-2. The connector bead corresponding to nbd1 and nbd2 are named CB1 and CB2 respectively, while the paratopes are referred to as P1 and P2 respectively (see Fig.3). In addition to the various bonded and non-bonded interactions described in the previous section, the angles L10-CB1-P1, P2-CB2-L1, L2-L1-CB2 and L9-L10-CB1 were restrained to 180° using a harmonic angle bending potential of the form of eq. (2) with *θ*_0_ = 180° employing stiffer bending coefficients as compared to the angles corresponding to the linker. On top of this, for the SBCG systems, the nanobodies were restrained from rotating about their respective long axes by restraining a dihedral formed by L10-CB1 (L1-CB2) and two beads of nbd-1 (nbd-2) to their starting values throughout the simulation. The extra restraints were introduced to mimic the fact that, in real-life systems, the bonds between the linker and the nanobody restrict the angular motion of the nanobody about the CB1/2-P1/2 axis.

### PMF calculation

The PMF calculations have been performed using the umbrella sampling (US) technique along two different reaction coordinates (RCs). The two RCs are named *ρ*_*z*−*proj*_ and *ρ*_*x*−*y*_. The former is the projection of the vector joining the tethering point to the center of mass of nbd-1 on the *z*-axis (see Fig.4 (A)). The latter is the projection of the same vector on the *x* −*y* plane with the condition that *ρ*_*z*−*proj*_ = 5 *σ*, which represents a condition where nbd-1 is close to the wall to which nbd-2 is tethered (see Fig.4 (B)). The RC values varied from 3 to 30, 40, 50, 60 and 70 *σ* for the 10-, 20- and 30-, 40- and 50-mer linker systems, respectively, with windows at gaps of 0.5 *σ*, leading to 55, 75, 95, 115 and 135 simulation windows for the SPH systems. For the SBCG systems, the RC values varied from 5 to 35, 45 and 55 *σ* for the 10-, 20- and 30-mer linker systems, leading to 61, 81 and 101 windows. The US simulations in different windows were performed in parallel. To generate the starting structure for each window, a short simulation was performed with the free nanobody being dragged from the initial to the final value of the reaction coordinate with an equilibration time of 5 × 10^5^ steps at each value, and the final snapshots at each value were used as the starting structures for different windows. Starting from the structures thus generated, simulations lasting for 5 × 10^7^ time steps were performed in each US window.

### Calculation of flight/residence times

The different kinds of simulations performed to calculate the flight and residence times are listed in Table I. We simulated a system consisting of a group of 25 diabodies arranged in a 5 × 5 square lattice, tethered to the lower *Z* surface (see Fig. 2 of SI (40)). The distance between two neighboring diabodies was set such that no interaction would be possible between them at any time. An initial simulation of 15 *µ*s was performed. The final structure was used to perform three independent simulations lasting for 30 *µ*s for the SPH system (linker lengths: 10−60). For the SBCG system (linker lengths: 10−30) one single simulation lasting 15 *µ*s was performed for each linker length for comparison. In these simulations, the *Z*-walls were repulsive. Additional simulations were performed for the SPH system for all different linker lengths, with slightly attractive *z*-walls. The value of *ϵ* for these simulations was set to 1.5 *k*_*B*_*T* and 2.5 *k*_*B*_*T* for two different sets of simulations. The *z*-coordinate of P1 was recorded and a threshold *z*_*th*_ = 3.5 *σ* between P1 and the tethering wall was set to distinguish between flight and residence. More precisely, a series of consecutive simulation frames during which P1 stayed below the threshold was considered to be a residence event, while flight events (and corresponding times) were associated with consecutive frames where the paratope remained above the threshold. Taking an average over the 25 nanobodies (nbd-1) and three independent simulations, we computed the average flight and residence times and also calculated the corresponding distributions.

## RESULTS AND DISCUSSION

We performed a 10 ns long simulation of the SBCG nanobody and used the trajectory to measure the radius of gyration of the coarse-grain model. We found the average over the trajectory to be 3.99 ±0.06 *σ*. The same calculation over the last 10 ns of the atomistic trajectory of the 1qd0 structure yielded an average value of 4.12 ± 0.02 *σ*, which confirmed the soundness of our SBCG-based approach.

### Free energy profiles

The PMF profiles as a function of *ρ*_*z*−*proj*_ are shown in Fig. 5 (A) and (C) for the SPH and SBCG diabodies, respectively. In spite of the PMF profiles having similar shape, there are a few notable differences. For the 10-mer system, for example, the PMF profile is flat in 15*σ* ≤ *ρ*_*z*−*proj*_ ≤ 25*σ* for the SBCG system, while the same region for the SPH system occurs at shorter distances, extending between 8*σ* and 20*σ*. In addition, for small values of RC, the profiles for the SBCG system have a larger slope as compared to the SPH system, which means that the SBCG nanobody faces a larger (entropic) repulsive force from the fixed wall than the SPH nanobody. The derivative of the PMF profiles is shown in Fig. 3 of SI (40). These differences suggest that the different coarse-graining schemes highlight differences in the dynamics near the tethering wall. In particular, an excessively simple spherical model seems to fall short of capturing important features of the interactions with a boundary.

**Figure 3:**
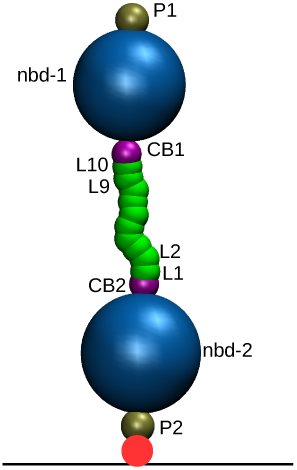
A 10-mer SPH diabody with the labels of different beads. The same naming scheme is used for the SBCG diabody. The tethering point is represented by the red bead.

**Figure 4:**
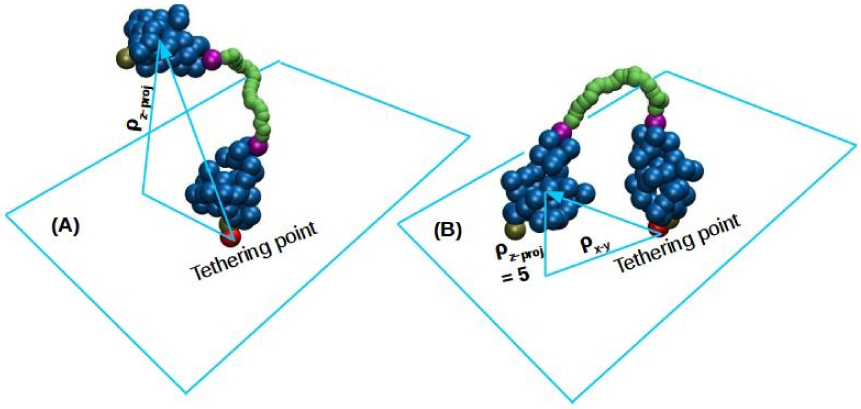
Reaction coordinates. Schematic representation of (A) *ρ*_*z*− *proj*_ and (B) *ρ*_*x*− *y*_. The red bead is the tethering point.

**Figure 5:**
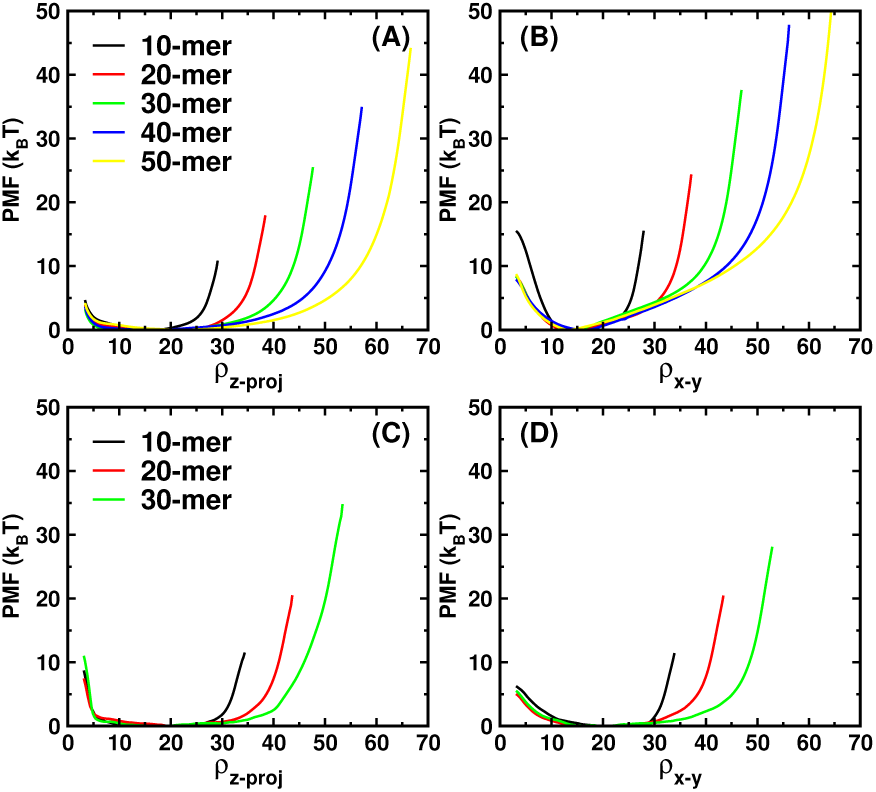
PMF profiles for tethered diabodies. PMF profiles as a function of the *z*-projection of the vector joining the tethering point to the center of mass of the free nanobody for the (A) SPH and (C) SBCG systems. PMF profiles as a function of the projection on the *xy* plane of the vector joining the tethering point to the center of mass of the free nanobody, with the free NB restrained to stay at a *z*-projection of 5 *σ* for the (B) SPH and (D) SBCG systems. The *x*-axis is represented in units of *σ*.

The PMFs are portrayed as a function of *ρ*_*x*−*y*_ in Figs. 5 (B) and (D). The profiles for the two models appear very different. While the PMFs for the SPH systems increase steadily in a linear fashion after *ρ*_*x*−*y*_ ∼ 15 *σ*, the profiles for the SBCG systems are flatter and start rising at a much later stage. Fig. 6 (A) shows the average of all values of the (normalized) *ρ*_*x*−*y*_ values for which PMF(*ρ*_*x*−*y*_) ≤ *k*_*B*_*T* for *ρ*_*z*−*proj*_ = 5 *σ* as a function of the linker length. The error bars gauge the flatness of the PMF profiles, which is seen to vary markedly in the two models. More precisely, the PMF profiles are flat (in the *xy* plane) for short linkers, while the flatness decreases with increasing linker length. This indicates that a network of nanobodies linked with short linkers may be more effective at inter-epitope binding even for configurations with a large standard deviation in the inter-epitope distances.

**Figure 6:**
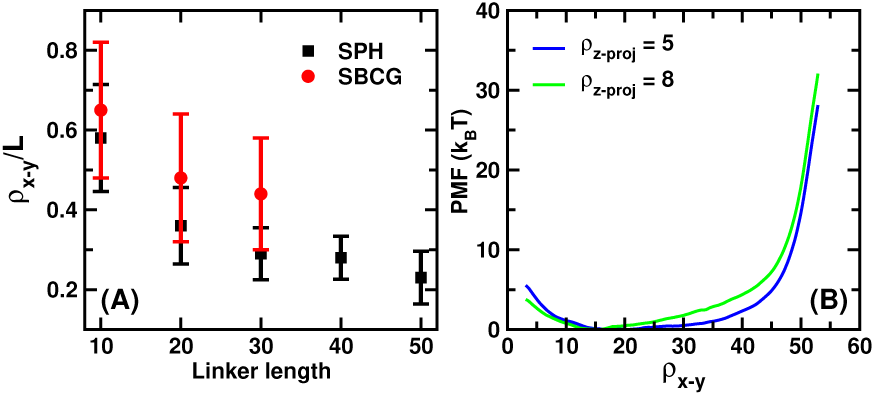
(A) Average of *ρ*_*x*−*y*_*/L* over the flat region of the PMF profiles as a function of linker length. The error bars quantify the extent of the flat region. (B) PMF profiles for the SBCG diabodies with the 30-mer linker as a function of *ρ*_*x*−*y*_ for two different values of *ρ*_*z*−*proj*_. The PMF profile for a wall-touching situation is different from the one where nbd-1 is farther from the fixed wall. In that case, the PMF profile for the SBCG diabody resembles that of the SPH diabody.

To better understand the differences displayed by the two models, we computed the PMF along *ρ*_*x*−*y*_ for different values of *ρ*_*z*−*proj*_ for the SBCG diabodies with 30-mer linkers. Fig. 6 (B) shows the profiles obtained for *ρ*_*z*−*proj*_ = 5 *σ* (nbd-1 in contact with the fixed wall) and *ρ*_*z*−*proj*_ = 8 *σ* (nbd-1 separated from the wall). For the larger value of *ρ*_*z*−*proj*_, the PMF profile appears similar to that of the SPH diabody, which indicates that the presence of the wall shapes the PMF profiles of the non-spherical model. We infer that for epitope systems with targets present very close to the cell-membrane, which acts as a soft wall, one might have a larger chance of inter-epitope binding as the wall seems to have a flattening effect on the SBCG PMF profiles. Overall, the differences between the PMF profiles of the two systems demonstrates that molecular shape may play a big role in controlling the interaction with the tethering wall.

One notices a steeper increase of the PMF as a function of *ρ*_*x*−*y*_ at low distances for the 10-mer as compared to other linker lengths. This effect is much more pronounced for the spherical molecules (Figs. 5 (B) and (D)). This indicates that when the linker length is comparable to the dimensions of the nanobodies, it is difficult to approach the epitopes close-by to the one to which the diabody is tethered. It is to be stressed that the slope is larger in case of the SPH system as compared to the SBCG system, which indicates that the shape of the nanobody is expected to play a role when it comes to inter-epitope binding for a high target density, especially for low linker lengths.

In the spirit of computationally aided molecular design, the PMF profiles as a function of *ρ*_*x*−*y*_ can be used to estimate the length of the linker required to efficiently result in multi-epitope binding for a given target geometry. With a knowledge of the epitope size and average inter-epitope distances, one can estimate, with a knowledge of the position of minima of the PMF profiles and also their degree of flatness, what length (or range of lengths) of the linker polymer would result in avidity. Simulations with bending modulus of the linker matching different polymers used as linkers in practical situations, like various peptides or nucleic acids, can help predict the appropriate stiffness of the linker for a given geometrical arrangement of epitopes. Tumor receptors like HER2 (43) have one epitope for natural antibodies while some engineered triple helix proteins called affibodies (44, 45) are known to engage a different region on the opposite side (46). With PMF calculations and knowledge of the position of minima as a function of a suitable reaction coordinates, one can predict, using MD simulation, the linker length that would maximize bispecific binding.

### *z*-distribution of the free paratope: comparison with models

An important observable that is tightly related to the statistics of flight and residence times is the distribution of the *z*-coordinate of the paratope of the free nanobody (bead P1). We computed these distributions for all the systems from the simulations performed to calculate the flight and residence times (see methods). It is interesting to compare the data for the SPH and SBCG systems with a simple model. The expression for the normalized equilibrium *z*-distribution of the free end of a Gaussian polymer, tethered at a height *z*_0_ from a reflecting wall, reads (47)

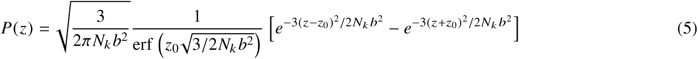

where *N*_*k*_ is the number of Kuhn monomers, *b* is the Kuhn length and *z*_0_ is the *z*-coordinate of the tethering point measured from the wall. In the limit *z*_0_ → 0, the expression reduces to (48):

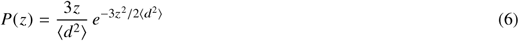

where we have introduced explicitly the average square end-to-end distance of the free polymer, ⟨*d*^2^ ⟩ = *N*_*k*_ *b*^2^.

It is interesting to inquire whether the data for the composite two-NB models can be described by an *effective* polymer model. The most obvious choice would be a Gaussian polymer with the same flexibility as the diabody linker, contour length equal to the contour length of the diabody measured from CB2 to the free paratope (P1) and tethered at the average height of CB2 above the lower *z* wall. A reasonable guess for the length of the effective model is thus *N*_*k*_ = (*N* + 2*r*)*/b*, where *N* is the number of monomers in the linker, *r* is half the distance between P1 and CB1 and *b* is the Kuhn length of the linker polymer (see again Fig. **??**). We set *z*_0_ = 7.5 *σ*, which is the average height for CB2 measured from the wall for the SPH system (see Fig. 5 of SI (40)). This would represent a polymer tethered at *z*_0_ = 7.5 *σ* distance units above the tethering wall and having a contour length equal to the linker and nbd1 combined. Our linker has a persistence length of ℓ_*p*_ ≃ 1.5 *σ* (see SI), which leads to a Kuhn length *b* = 2ℓ_*p*_ = 3 *σ*.

Fig.7 shows the comparison of the effective model with the data for the SPH and the SBCG systems. It is apparent that the distribution for the diabodies describes a higher representation of large *z* values and a reduced representation of small *z* values when compared to the tethered polymer. This is an expected consequence of the higher entropic repulsive force exerted by the wall on the free bead, a mechanism akin to the entropic pulling force demonstrated in the disassembling and translocating action of Hsp70s chaperones (49). More precisely, the conformational space near the wall is restricted more severely for the diabodies, due to presence of the nanobodies at the two ends of the polymer, than for a *bare* polymer with the same contour length and flexibility. The extent of the difference between the simulation and the model suggests extended, composite molecules such as our diabodies, belong to a different universality class altogether. One can, however, use eq. (5) or (6) to determine the effective Kuhn length 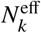 of an effective polymer with the same persistence length ℓ_*p*_ as the linker in the diabodies. For this, we fit the distributions of the 40-, 50- and 60-mer SPH systems with eq. (6) (see Fig.6 of SI) with *b* = 3 *σ*, which leads to 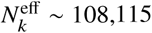 and 119 for 40-, 50- and 60-mer diabodies respectively.

**Figure 7:**
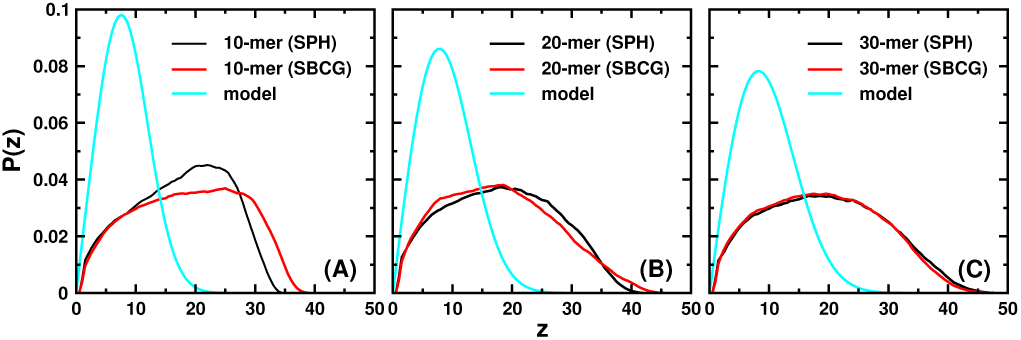
*z*-distribution of the free paratope (bead P1) for (A) 10- (B) 20- and (C) 30-mer SPH and SBCG systems plotted along with the equivalent Gaussian polymer approximation (5) (see text).

It is interesting to note that the distributions for the SBCG and the SPH systems differ to a highest extent when the linker length matches the dimensions of the linked NBs, i.e. for the shortest linker (10-mer), while they approach each other rapidly as the linker length increases and almost overlap for the 30-mer linker. This means that, as the statistics of the vertical coordinate above the wall is concerned, for linkers longer than approximately the size of the NBs, an equivalent SPH model can be used, entailing considerable simplification of the simulations.

### Flight/residence times: quantifying the kinetics of the second binding

The statistics of the flight/residence times (F/RT) of the free NB are crucial observables, as they embody the kinetic determinants of the second binding, hence can help quantify avidity. The F/RTs are defined as stretches of consecutive frames that the paratope of the free NB (bead P1) spends above/below, respectively, a fixed threshold height *z*_*th*_ from the tethering wall. Let us denote with *P*_*f*_ (*t*) and *P*_*r*_ (*t*) the equilibrium distributions of flight and residence times. The corresponding complementary cumulative distributions, *S*_*f*_ (*t*) and *S*_*r*_ (*t*), defined as

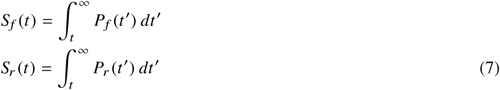

coincide with the survival probabilities relating to the corresponding domains, 𝒟_*f*_ ={*z* ∈ [0, *L*] | *z* ≥ *z*_*th*_} and 𝒟_*r*_ = {*z* ∈ [0, *L*] | *z* < *z*_*th*_, *L* being the side of the simulation box along the *z* direction. We note that the domain _*r*_ can also be regarded as the paratope-wall *interaction* domain. More precisely, *S*_*f*_ (*t*) represents the fraction of flight events whose duration exceeds *t*, hence this is nothing but the probability that the paratope be still in region 𝒟_*f*_ after a time *t*, i.e. indeed the survival probability for domain 𝒟_*f*_. Analogously, *S*_*r*_ (*t*) represents the survival probability for domain 𝒟_*r*_. The corresponding distribution of exit times (in the sense of first passage times), 𝒫_*f*_ (*t*) and 𝒫_*r*_ (*t*), can then be computed straightforwardly as (50)

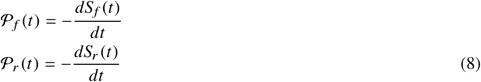

The inverse mean-exit times can be considered as measures of the escape rate from the corresponding domains. Therefore, combining eqs. (7) and (8), it is possible to estimate the on and off rates directly from the series of flight and residence times, as

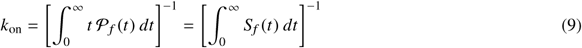

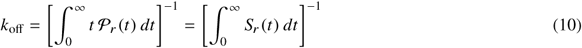

To calculate the flight (*t* _*f*_) and residence times (*t*_*r*_) numerically, the *z*-coordinate of the paratope was monitored through the simulation. A threshold *z*-height of *z*_*th*_ = 3.5 *σ* was set to define whether the paratope is in the flight or residence regions. If the paratope stayed above or below the threshold for at least 0.5 ns, the corresponding trajectory stretch was designated to correspond to a flight or residence event, respectively. This additional constraint was tailored specifically to avoid short recrossing events that could bias the statistics unphysically at short times. Data for such events were accumulated over 30 *µ*s-long trajectories of 25 non-interacting tethered diabodies. A set of 3 such simulations were performed for each linker length. In addition, the simulations were performed for both repulsive and attractive tethering walls (with attractive energies equal to 1.5 *k*_*B*_*T* and 2.5 *k*_*B*_*T*), with the aim of assessing the effect on the paratope-wall kinetics of some weak non-specific attraction between the protein and the wall/membrane.

Fig. 8 (upper panels) illustrates the calculation of the on and off rates as described by formulas (9) and (10). The probability per unit time that the paratope enter the interaction domain appears of the order of tens of *µ*s^−1^, while the probability per unit time that it exit the same domain turns out to be about ten times higher. Interestingly, the SBCG model shows a higher on-rate than the spherical model (with a pure repulsive wall). At the same time, the exit probability is higher for the SBCG model.

**Figure 8:**
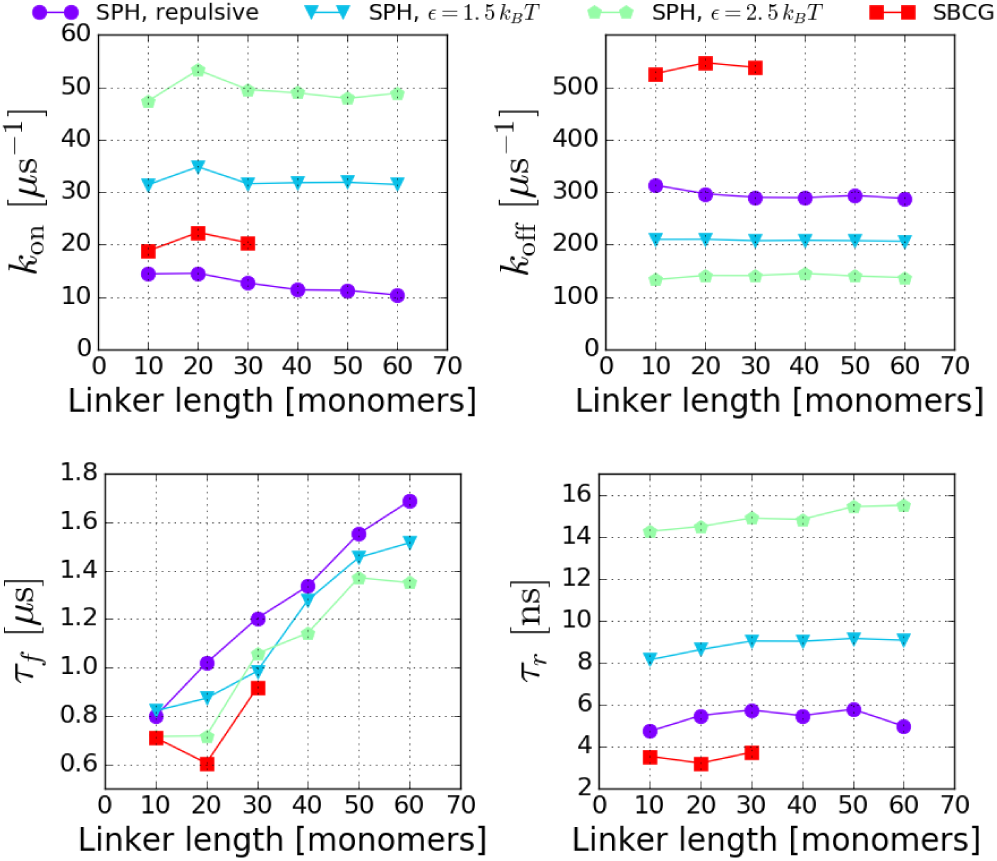
Upper panels. On and off rates describing the kinetics of paratope entering and exiting the interaction domain 𝒟_*r*_ (i.e. *z* < *z*_*th*_), computed according to the prescriptions (9) and (10). Bottom panels. Time constants of the exponential tails of the survival probabilities. These can be thought of as the average time of the rare, very long events. The data for the SPH model refer to the simulation set #2 (see Table I)

Fig. 8 also demonstrates that a weak non-specific attraction between the NBs and the wall of the order *ϵ* ≃ 1 − 2 *k*_*B*_*T* increases the on-rate (i.e. decreases the overall survival probability in the flight domain) and decreases the off-rate (i.e. makes journeys of the paratope in the interaction domains longer). More specifically, these data refer to a modified SPH-wall system with non-specific isotropic attraction (LJ) between the wall and nbd-1/2 and P1/2. It is interesting to observe that the gain afforded by a weak attractive wall in terms of on-rate, as gauged by *k*_on_ (*ϵ, N*)*/k*_on_ (0, *N*), is found to increase with the linker length *N*. In fact, while, *k*_on_ (0, *N*) decreases with *N*, a weak attraction makes *k*_on_ (*ϵ, N*) nearly insensitive to variations in the linker length. This possibly reflects the fact that the entropic cost associated with entering the interaction domain decreases with increasing linker length in the presence of attraction between the paratope/NB system and the wall. Conversely, the ratio *k*_off_ (*ϵ, N*)*/k*_off_ (0, *N*) appears to remain constant as *N* increases.

It is instructive to inspect in more detail the survival probabilities. Fig. 9 depicts the survival curves for both the flight and paratope-wall interaction domain. The short time behavior (*t* ≲;S 0.5 *µ*s) appears to follow an inverse power law with exponent 1*/*2 (see dashed lines in Fig. 9), irrespective of the linker length. This is the expected behaviour for the survival probability of a free random walk in three dimensions (50). This means that short survival times in either domain are dominated by the unconstrained diffusion of the paratope. By contrast, longer survival events depend markedly on the length of the linker and are distributed exponentially. The inset in Fig. 9 makes this point very clear in the case of the function *S*_*f*_ (*t*) for the SPH model. We find that this is a general feature of the tails of the paratope survival probabilities in either region. As it shows from the figures, the tails of the flight times depend on the linker length, while those of the residence times much less so. In order to have some insight into the tail, rare-event region, we might reason as follows.

**Figure 9:**
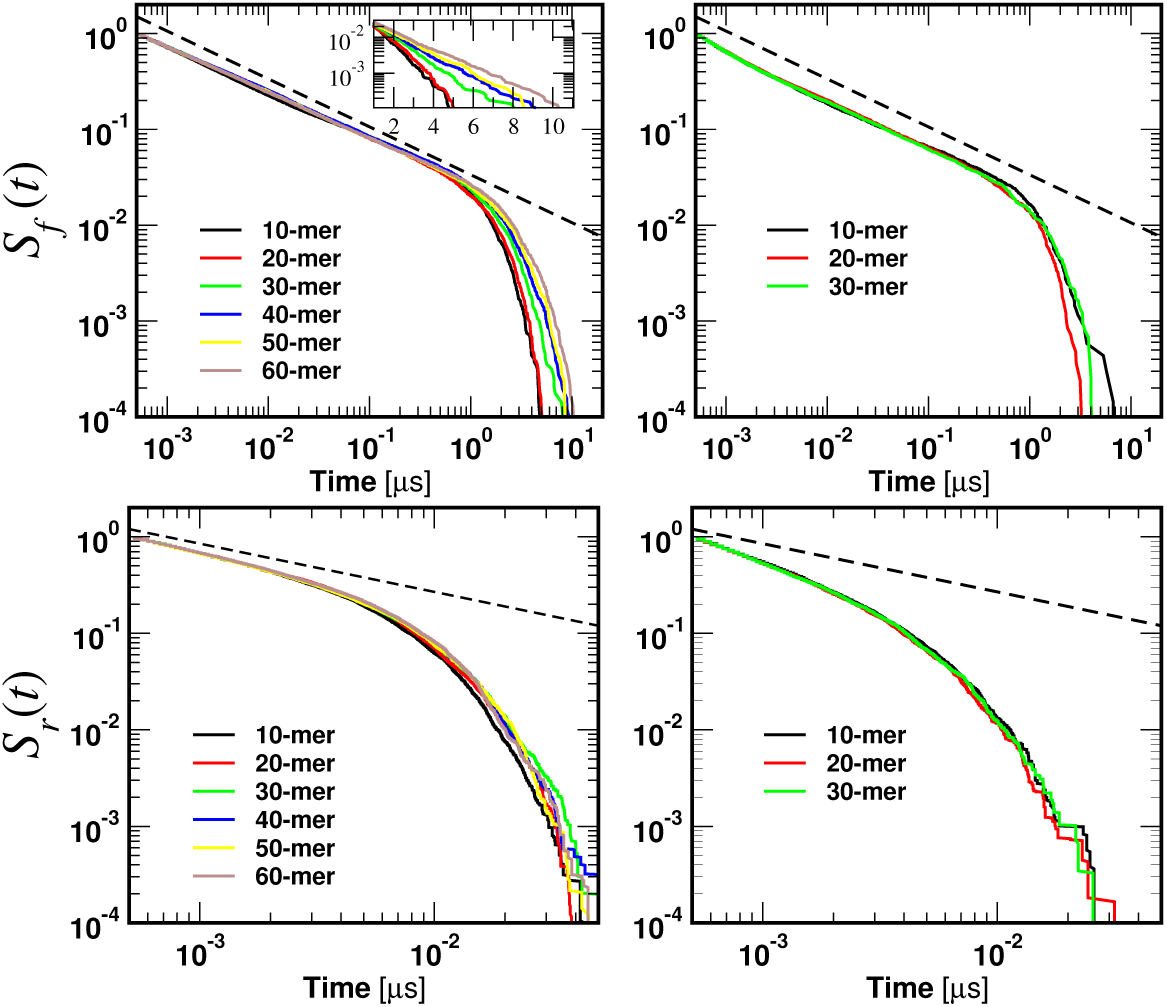
Survival probability of the epitope in the flight (*z* ≥ *z*_*th*_, top) and residence (*z* < *z*_*th*_, bottom) domains for (left) the SPH system and (right) SBCG system for different linker lengths. The dashed lines are plots of power laws of the kind *t*^−1/2^. The inset shows a close-up in lin-log scale of the tails of the flight survival probability, which makes the exponential decay clearly visible. The plots for the attractive walls are provided in Fig. 7 of SI (40).

We can safely assume that rare events make uncorrelated time series. In this case, the statistics of return events will be specified by the configurational probability (independent of time) that the paratope be in the relevant regions, either *z* ≥ *z*_*th*_ or *z* < *z*_*th*_. In turn, this will depend on the conformational statistics of the NB-polymer systems and, in the absence of an appropriate analytical model, can be easily determined from our PMF calculations (see Fig. 5). Let us denote with *P*_>_, *P*_<_ the equilibrium probability that the free NB be above or below the threshold *z*_*th*_, respectively. In this case, the probability *P*_*a*_ (*k*) and *P*_*b*_ (*k*) of observing *k* consecutive sampled frames above (a) or below (b) *z*_*th*_, respectively, can be computed as

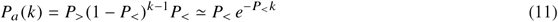

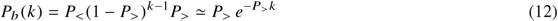

If for the sake of the argument we take the Gaussian tethered model (6) as a reference case, it is readily seen that

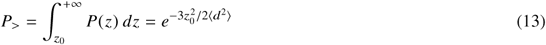

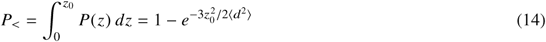

If we introduce the time decay constants *τ*_*f*_, *τ*_*r*_ of the exponential tails of the survival probabilities in the flight and interaction domains, respectively, Eqs. (13) and (14) entail

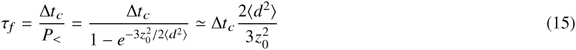

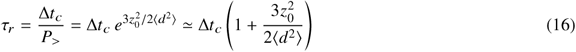

where Δ*t*_*c*_ is a time of the order of the typical correlation time of consecutive frames and in the last passages we have made use of the fact that 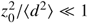.

Fig. 8 indeed shows that *τ*_*f*_ increases linearly with *N* (⟨*d*^2^⟩ ∝ *N*) for the spherical model, according to the prediction (15). It is interesting to observe that the slope does not seem to depend on the value of the weak attractive energy *ϵ*. This is expected, as rare, long flight events are dominated by the statistical weight of the configurations of the combined NB/linker molecule away from the wall. The simple calculation leading to eq. (16) also correctly explains the observed reduced variability of residence times as the linker length is increased (see again Fig. 8, bottom right panel). However, a closer inspection reveals that the average duration of rare, long residence events increases with the linker length *N* in the presence of an attractive interaction between the paratope/NB system and the wall, even though with a much smaller slope than for the increase of *τ*_*f*_. Overall, we conclude that the effect of a weak non-specific interaction with the wall decreases the average duration of rare, long flights and residence events. It is interesting to observe that the SBCG model (with a repulsive wall) displays the shortest duration of rare long flights (Fig. 8, bottom left), even shorter than for the spherical model in the presence of the most attractive wall (*ϵ* = 2.5*k*_*B*_*T*). Moreover, there seems to be an optimum (a pronounced dip) at a linker length that is approximately the same size of the attached NBs (*N* = 20).

While rare, long flight and residence times are on the *µ*s and tens of ns scales (exponential tails), respectively, the average values ⟨*t* _*f*_⟩, ⟨*t*_*r*_⟩ are dominated by the short-time power-law behaviour. Fig. 10 reveals that average flight times turn out to be of the order of about 10^2^ ns, while residence times are about 50 times shorter, of the order of 2 − 3 ns. Furthermore, one can appreciate that the SBCG model systematically displays shorter flight and residence times with respect to the spherical model. As for rare events, this feature should be attributed to the shape and intrinsic flexibility of the SBCG NBs as compared to equivalent rigid spheres of the same size.

**Figure 10:**
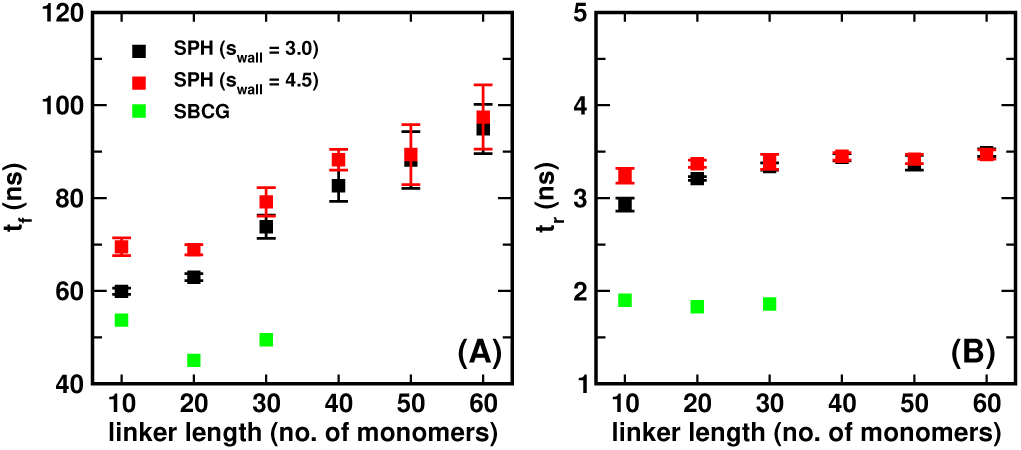
Average flight times (A) and average residence times (B), computed over the trajectories of 25 non-interacting, tethered diabodies. A distance cutoff *z*_0_ = 3.5 *σ* between the tethering wall and the free paratope (P1) was used to calculate the values. A jump above (a plunge below) the cutoff was considered to be a flight (residence) event only if it lasted for more than 0.5 ns, The systems compared comprise two different variants of the SPH model, as described in table 1, and the SBCG system.

**Figure 11:**
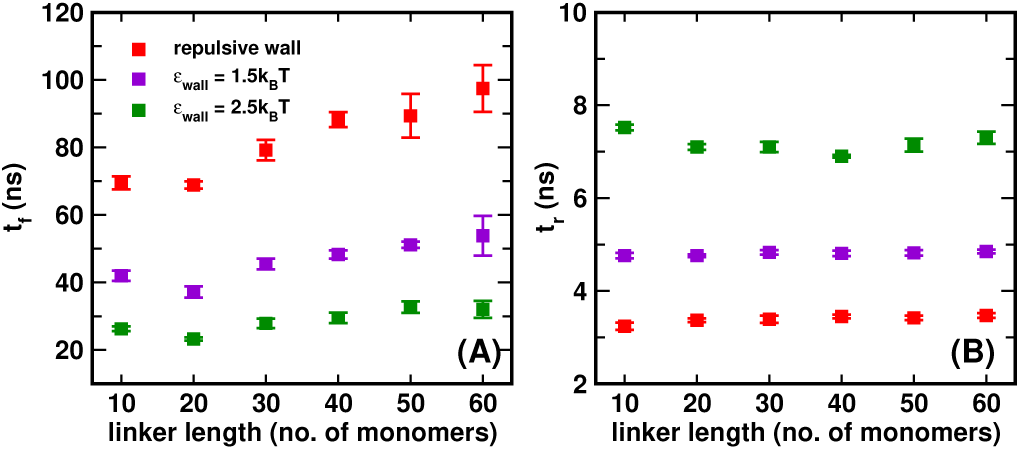
Repulsive vs. attractive wall. Comparison of (A) average flight times and (B) average residence times for SPH systems for repulsive and attractive tethering walls computed as described in Fig. 10.

It is interesting to observe that shorter linker (10-mers) correspond to rather unfavorable situations (large flight times). Remarkably, increasing the linker length from *N* = 10 results in a substantial reduction in the duration of the flight stretches, an effect that is more pronounced for the SBCG model (see again Fig. 10 A). A close inspection of the trajectories shows that for the 10-mer linker, for which the linker length is less than the size of the nanobodies, the tethered unit, due to its steric extension, introduces a steep entropic barrier for the free NB when it approaches the wall (see Fig.8 of SI (40)). The consequence of such steric repulsion and its more pronounced effects in the case of the 10-mer linkers can also be seen from Fig. 7, where the peak of the *z*-distributions of the free paratope (bead P1) are farthest away from that of the simple equivalent Gaussian polymer for the 10-mer, and more so for the SBCG diabody. This effect is present for the SPH model too, albeit less marked. Again, also for average values, there seems to be an *optimum* length of the linker that minimizes the average time spent by the free NBs away from the tethering surface, that approximately matches the size of the binding units themselves. All the data for *t* _*f*_ and *t*_*r*_ for this set of simulations are reported in tabulated form in tables I to V in the SI.

In order to investigate further the details of the NB-wall interactions, we performed the simulations with repulsive tethering walls for two different values of the LJ repulsive length *s*_*wall*_ (4.5 *σ* and 3.0 *σ*) for nbd-2 (spherical model). We observe that for smaller linker lengths (10 − 30-mers), flight times are shorter for a smaller value of *s*_*wall*_. The effect is negligible for the longer linkers, and seems, in fact, inversely proportional to the linker length. The effect arises because the value of *s*_*wall*_ determines how nbd-2 (the tethered NB) would interact with the tethering wall and how much it is able to bend. This is expected to affect the value of ⟨*t* _*f*_⟩, more so in presence of the external harmonic potential that restrains the value of the angle between P2-CB2-L1 (the axis of the tethered sphere) to 180°. One would then expect that how much nbd-2 can bend would depend on the shape of the nanobody, as the shape would then determine the dynamics and effective interaction of the nanobody with the tethering wall.

The bending propensity of the tethered unit can be gauged by calculating the height distribution of CB2 (see fig. 3) measured from the tethering wall surface for the SPH and the SBCG systems (see Fig.5 of SI (40)). From the plot we see that the very low heights have a significant probability in case of the SBCG diabodies, which may contribute to the lower values of ⟨*t* _*f*_⟩. By contrast, for the SPH diabodies there is an abrupt lower cutoff depending on the value of *s*_*wall*_. From Fig. 10 we notice that the values of ⟨*t* _*f*_⟩ for the SBCG diabodies are substantially shorter as compared to the SPH diabodies. For the 30-mer linker, for example, the SPH system has ⟨*t* _*f*_⟩ ∼ 80 ns while for the SBCG system ⟨*t* _*f*_⟩ ∼ 50 ns. It thus seems that it is not an obvious task to reproduce the interaction with a wall within by a model that preserves the shape of the atomistic structure of the NB via a simple spherical representation. Thus the SBCG model seems more appropriate to calculate relevant dynamical parameters, such as the on-rate for second binding, that are expected to rely substantially on the details of the interactions with a wall/membrane.

A closer look at the average residence times reveals that, while ⟨*t*_*r*_⟩ shows negligible dependence on the linker length, the dependence on the model is noticeable. The residence time for the SPH diabodies is ∼ 3.5 ns, while the values for the SBCG diabodies is close to 2 ns. The difference can be attributed to the fact that the SBCG nanobody likely generates larger reaction forces from the tethering walls on behalf of its fluctuating structure (see Fig.3 of SI (40)), thus reducing the time it stays near the wall. By contrast, the SPH nanobody would face a smaller reaction from the wall, given its rigid and fixed surface. Here again one can appreciate the importance of the model being used.

Finally, to ascertain that the length of the flight-residence times simulations was sufficient to arrive at converged values of ⟨*t* _*f*_⟩ and ⟨*t*_*r*_⟩, we performed the calculation for different durations of the simulations, and checked how the calculated values changed as a function of the simulation length. For the 30-mer linker, for example, ⟨*t* _*f*_⟩ started from a value of 85 ± 22 ns for a simulation length of 3.75 *µ*s and slowly converged to the reported value of ∼ 77 ns for a simulation length of 15 *µ*s and stayed close to that for longer simulation lengths, suggesting that our simulation length of 30 *µ*s is appropriate for producing converged results. The values of ⟨*t* _*f*_⟩ as a function of simulation length for the SPH systems with *N* = 30 and *N* = 60 linkers are shown in Fig.9 of SI (40).

## CONCLUSIONS AND PERSPECTIVES

In this work we have introduced two different coarse-grained models of polymer-linked, two-nanobody molecules, as a simple but paradigmatic example of novel immunotherapy agents that are increasingly being developed in a variety of contexts. More precisely, while the linker has been modeled invariably as a bead-and-spring system (stretching and bending), the nanobodies have been represened either as single large rigid spheres, or as collections of small spheres suitably connected by springs. Such representation was parameterized in a bottom-up philosophy directly from atomistic simulations in explicit solvent. The latter scheme led to binding units displaying the same *shape* and large-scale flexibility as the atomistic systems.

The aim of this work was to lay the bases of coarse-grained, computationally aided drug design in this area from the firm standpoint of statistical and computational physics. In the spirit of the accepted model of sequential binding of bivalent molecules (51), whereby bivalent agents first bind to a target-covered surface from the bulk, and subsequently dynamically explore the surroundings for a second target within reach (either on the same or on a facing surface), our main focus was to elucidate the physics of the latter kinetic step. For this purpose, we have mainly focussed on the kinetic and equilibrium properties of a molecule tethered to a wall through one of its binding units (NB), in order to investigate the main dynamical and structural determinants of the second binding. In particular, we aimed at investigating (i) to what extent the degree of coarse-graining may impact the dynamics of the combined linker/free NB system and (ii) the interaction dynamics of the free paratope (the active binding site carried by the NB) with the surface.

In the first part of this work, we have shown that the calculation of potentials of mean force (PMF) along specific, one-dimensional collective coordinates can provide considerable insight into the nature, strength and range of the thermodynamics forces that govern the motion of the free paratope. Furthermore, such calculations may constitute a precious tool to investigate the role of such forces in the dynamics of the paratope-wall encounter, which, in turn, governs the kinetics of the second binding.

We aim at illustrating this aspect in a forthcoming publication. For example, the PMF can be fruitfully used as an effective potential in approaches based on the Smoluchowski equation or on first-passage processes (50).

In the second part of the paper, we have delved into the kinetics of the paratope-wall encounters. To this end, we have developed a general strategy based on dissecting an equilibrium trajectory of the free paratope in flight and residence stretches, depending on whether the active site on the free NB was found above or below, respectively, a critical interaction threshold close to the wall in the vertical direction. We have shown that the encounter and escape kinetics with respect to the wall are simply related to the survival probability of the paratope in the flight and interaction domains, respectively, which can be simply computed from the series of flight and residence times observed over a long MD trajectory. We have illustrated how this method allows one to estimate the kinetic constants of the second binding, *k*_on_ and *k*_off_, in the presence of a purely repulsive wall and with weak, non-specific interactions between the free NB and the wall. Our simple method not only allows one to quantify the role of factors such as the linker length and flexibility (not considered in this study) and non-specific interactions on the average flight/residence times. It also makes clear and quantify the role and weight of rare, long-duration events that show up in the exponential tails of the survival probabilities.

It is worth stressing that the statistics of rare events is by no means a secondary issue in this context, as in many real-life situations the binding kinetics of such molecules is expected to be dominated by fluctuations, e.g. due to low copy numbers or tiny reaction volumes. For example, this is the case of novel bivalent and bispecific diabodies engineered to bind within the immune synapse, i.e. at the interface of two facing membranes, on the effector cell (NK or B-cell) on one side and on epitopes on a tumor cell on the other side. The synapse covering an area of the order of 100 *µ*m^2^ for a cell-cell separation of about 15 nm (24, 25), the role of fluctuations in the number of bridging molecules is expected to be important, which likely makes the statistics of rare events a major determinant of the binding kinetics.

## AUTHOR CONTRIBUTIONS

FP conceived the study. SS performed the simulations. SS and FP analyzed the data and wrote the paper.

## ACKNOWLEDGEMENT

The authors acknowledge the French ministry of higher education under the PIA2 project BIOS for funding.

## REFERENCES

1. A Desmyter, S Spinelli, A Roussel, C Cambillau, Current Opinion in Structural Biology. 32, 1–8 (2015).

2. P Chames, M V Regenmortel, E Weiss, D Baty, Br J Pharmacol. 157(2), 220–233 (2009).

3. G Behar, S Sibéril, A Groulet, P Chames, M Pugnière, C Boix, C Sautès-Fridman, J L Teillaud, D Baty, Protein Engineering Design and Selection 21(1), 1–10 (2008).

4. M Turini, P Chames, P Bruhns, D Baty, B Kerfele, Oncotarget. 5(14), 5304–5319 (2014).

5. P Bannas, A Lenz, V Kunick, W Fumey, B Rissiek, J Schmid, F Haag, A Leingärtner, M Trepel, G Adam and F Koch-Nolte, J Vis Exp 98, e52462 (2015).

6. P Bannas, A Lenz, V Kunick, L Well, W Fumey, B Rissiek, F Haag, J Schmid, K Schütze, A Eichhoff, M Trepel, G Adam, H Ittrich, F Koch-Nolte, Contrast Media Mol Imaging 10(5), 367–78 (2015).

7. M Kijanka, B Dorresteijn, S Oliveira, P M van Bergen en Henegouwen, Nanomedicine (Lond) 10(1), 161–74 (2015).

8. P Bannas, A Lenz, V Kunick, L Well, W Fumey, B Rissiek, F Haag, J Schmid, K Schütze, A Eichhoff, M Trepel, G Adam, H Ittrich and F Koch-Nolte, Contrast Media Mol Imaging. 2015 10(5), 367–78 (2015).

9. S A Kostelny, M S Cole and J Y Tso, J Immunol. 148(5), 1547–53 (1992).

10. C De Michele, P De Los Rios, G Foffi, F Piazza, PLOS Computational Biology, 12(3), e1004752 (2016).

11. C Kimchi-Sarfaty, T Schiller, N Hamasaki-Katagiri, M A Khan, C Yanover and Z E Sauna, 34(10), 534–548 (2013).

12. H A Lagassé, A Alexaki, V L Simhadri, N H Katagiri, W Jankowski, Z E Sauna and C Kimchi-Sarfaty. Recent advances in (therapeutic protein) drug development. F1000Res 7(6), 113 (2017). Biophysical Journal Template

13. E Chertova, J W Bess, B Crise Jr., R C Sowder II, T M Schaden, J M Hilburn, J A Hoxie, R E Benveniste, J D Lifson, L E Henderson and L O Arthur, J. Virol. 76, 5315–5325 (2002).

14. J Liu, A Bartesaghi, M J Borgnia, G Sapiro, and S Subramaniam, Nature 455, 109–113 (2008).

15. P Zhu, J Liu, J Bess Jr., E Chertova, J D Lifson, H Grisé, G A Ofek, K A Taylor, and K H Roux, Nature 441, 847–852 (2006).

16. J S Klein and P J Bjorkman. PLoS Pathog 6, e1000908 (2010).

17. H Mouquet, J F Scheid, M J Zoller, M Krogsgaard, R G Ott, S Shukair, M N Artyomov, J Pietzsch, M Connors, F Pereyra, B D Walker, D D Ho, P C Wilson, M S Seaman, H N Eisen, A K Chakraborty, T J Hope, J V Ravetch, H Wardemann, M C Nussenzweig, Nature 467, 591–595 (2010).

18. H Wu, D S Pfarr, Y Tang, L L An, N K Patel, J D Watkins, W D Huse, P A Kiener and J F Young, J. Mol. Biol. 350, 126–144 (2005).

19. R P Galimidi, J S Klein, A P West Jr. and P J Bjorkman, Cell 160, 433–446, a2015 Elsevier Inc (2015).

20. S Jähnichen, C Blanchetot, D Maussang, M Gonzalez-Pajuelo, K Y Chow, L Bosch, S De Vrieze, B Serruys, H Ulrichts, W Vandevelde, M Saunders, H J De Haard, D Schols, R Leurs, P Vanlandschoot, T Verrips, M J Smit, Proc Natl Acad Sci U S A 107(47), 20565–70 (2010).

21. J Zhang, J Tanha, T Hirama, N H Khieu, H Tong-Sevinc, E Stone, J R Brisson and C R MacKenzie. J Mol Biol. 335(1), 49–56 (2004).

22. T Yang, O K Baryshnikova, H Mao, M A Holden, P S Cremer, J. Am. Chem. Soc. 125(16), 4779–4784 (2003).

23. C Fasting, C A Schalley, M Weber, O Seitz, S Hecht, B Koksch, J Dernedde, C Graf, E Knapp, R Haag, Angewandte Chemie 51(42), 10472–98 (2012).

24. V L Schiavom P Robert, L Limozin, P Bongrand, PLoS One 7(9), e44070 (2012).

25. González C1, Chames P2, Kerfelec B2, Baty D2, Robert P3, Limozin L4, Biophys J 116(8), 1516–1526 (2019).

26. M van Rosmalen and M K Maarten, Biochemistry 56(50), 6565–6574 (2017).

27. C Xiaoying, Z Jennica and S Wei-Chiang, Adv Drug Deliv Rev 65(10), 1357–1369 (2013).

28. J S Klein, S Jiang, R P Galimidi, J R Keeffe and P J Bjorkman, Protein Engineering, Design and Selection, 27(10), 325–330 (2014).

29. R Arai, H Ueda, A Kitayama, N Kamiya and T Nagamune, Protein Engineering, Design and Selection, 14(8), 529–532 (2001).

30. H Zhao and A Caflisch, European Journal of Medicinal Chemistry 91 4e14 (2015).

31. J Jung, W Nishima, M Daniels, G Bascom, C Kobayashi, A Adedoyin, M Wall, A Lappala, D Phillips, W Fischer, C S Tung, T Schlick, Y Sugita, K Y Sanbonmatsu, Journal of Computational Chemistry 40(21), 1919–1930 (2019).

32. B Windisch, D Bray and T Duke, Biophys J 91(7), 2383–2392 (2006).

33. A Arkhipov, W H. Roos, G J L Wuite, and K Schulten, Biophys J 97(7), 2061–2069.

34. S Spinelli, L G Frenken, P Hermans, T Verrips, K Brown, M Tegoni and C Cambillau, Biochemistry 39(6), 1217–22 (2000).

35. M P Allen and D J Tildesley, Computer simulation of liquids, Oxford university press: New York, (1991).

36. J C Phillips, R Braun, W Wang, J Gumbart, E Tajkhorshid, E Villa, C Chipot, R D Skeel, L Kale and K Schulten, Journal of Computational Chemistry 26(16), 1781–1802 (2005).

37. R B Best, X Zhu, J Shim, P E M Lopes, J Mittal, M Feig, and A D MacKerell Jr., J. Chem. Theory Comput 8(9), 3257–3273 (2012).

38. W Humphrey, A Dalke and K Schulten, J Mol Graphics 14(1), 33–38 (1996).

39. J D Weeks, D Chandler, H C Andersen, The Journal of Chemical Physics, 54(12), 5237Ð5247 (1971).

40. See Supplemental Material at [URL] for the details of the system simulated for calculating the flight/residence times, comparison of the interaction force between the tethering wall and nbd-1 for the two models, the radial distribution of the position of P1, Distribution of the normalized height of CB2 from the tethering wall for the 10-mer SPH system (repulsive wall, σ_wall_ = 4.5 for nbd-2) and the 10-mer SBCG system, Steric repulsion between nbd-1 and nbd-2 for the 10-mer linker preventing nbd-1 from reaching close to the tethering wall, convergence of the flight time values as a function with simulation length and flight/residence times for different systems in tabulated form.

41. H Lee, R M Venable, A D MacKerell Jr., and R W Pastor, Biophys J. 95(4), 1590–1599 (2008).

42. S Plimpton, J Comp Phys 117, 1–19 (1995).

43. C Gutierrez and R Schiff, Arch Pathol Lab Med 135(1), 55–62 (2011).

44. F Y Frejd and K T Kim, Exp Mol Med, 49(3), (2017).

45. J Löfblom, J Feldwisch, V Tolmachev, J Carlsson, S Ståhl and F Y Frejd, FEBS Lett 584(12), 2670–80 (2010).

46. C Eigenbrot, M Ultsch, A Dubnovitsky, L Abrahmsén and T Härd, Proc Natl Acad Sci U S A 107(34), 15039–44 (2010).

47. K A Dill, S Bromberg, Molecular Driving Forces: Statistical Thermodynamics in Biology, Chemistry, Physics, and Nanoscience, 2nd ed., 2010 (New York: Garland Science (2010).

48. E A DiMarzio, J. Chem. Phys. 42, 2101 (1965).

49. P De Los Rios, A Ben-Zvi, O Slutsky, A Azem, P Goloubinoff, P. Proceedings of the National Academy of Sciences, 103(16), 6166Ð6171 (2006).

50. S Redner, A guide to First-Passage processes, Cambridge University Press, Cambridge (2001).

51. C De Michele, P De Los Rios, G Foffi, F Piazza, F. PLoS Computational Biology, 12(3):e1004752 (2016).

